# Arabidopsis *ABCC4* encodes a cytokinin efflux transporter and is involved in root system development

**DOI:** 10.1101/2024.05.14.594121

**Authors:** Takuya Uragami, Takatoshi Kiba, Mikiko Kojima, Yumiko Takebayashi, Yuzuru Tozawa, Yuki Hayashi, Toshinori Kinoshita, Hitoshi Sakakibara

**Affiliations:** Graduate School of Bioagricultural Sciences, Nagoya University, Chikusa, Nagoya 464-8601, Japan; RIKEN Center for Sustainable Resource Science, Tsurumi, Yokohama 230-0045, Japan; Graduate School of Science and Engineering, Saitama University, Sakura, Saitama 338-8570, Japan; Graduate School of Science and Institute of Transformative Bio-Molecules (WPI-ITbM), Nagoya University, Chikusa, Nagoya 464-8602, Japan

**Keywords:** ABC transporter, *Arabidopsis thaliana*, cytokinin, efflux transporter, root development

## Abstract

The directional and sequential flow of cytokinin in plants is organized by a complex network of transporters. Genes involved in several aspects of cytokinin transport have been characterized, but a large part of the elaborate system remains elusive. In this study, we have identified *ABCC4* as a cytokinin efflux transporter gene. Using a transient expression system in tobacco leaves, we screened Arabidopsis transporter genes and isolated *ATP-BINDING CASSETTE TRANSPORTER C4* (*ABCC4*). Further validation through drug-induced expression in Arabidopsis and heterologous expression in budding yeast revealed that ABCC4 effluxes the active form of cytokinins. During the seedling stage, *ABCC4* was highly expressed in roots, and its expression was up-regulated in response to cytokinin application. Loss-of-function mutants of *ABCC4* displayed enhanced primary root elongation, similar to mutants impaired in cytokinin biosynthesis or signaling, which was suppressed by exogenous *trans*-zeatin treatment. In contrast, overexpression of the gene led to suppression of root elongation. These results suggest that ABCC4 plays a role in the efflux of active cytokinin, thereby contributing to root growth regulation. Our findings contribute to unraveling the many complexities of cytokinin flow and enhance our understanding of the regulatory mechanisms underlying root system development in plants.

## Introduction

Cytokinins are a class of phytohormones involved in the regulation of various layers of plant growth and development, such as cell division, shoot development and regeneration, leaf senescence, and nutrient responses (Schaller et al., 2015; Kieber and Schaller, 2018; Cortleven et al., 2019; Wybouw and De Rybel, 2019; Sakakibara, 2021). Naturally occurring cytokinins possess a prenyl side chain attached to the *N^6^* position of adenine, giving rise to various forms such as *N^6^*-(Δ^2^-isopentenyl)-adenine (iP), *trans*-zeatin (tZ), and *cis*-zeatin (cZ), each having distinct side chain structures (Mok and Mok, 2001; Sakakibara, 2006). Differences in the side chain structures are closely linked to activity strength, as demonstrated by iP and tZ showing a higher affinity to their receptors compared to cZ across various plant species including *Arabidopsis thaliana* (Romanov et al., 2006; Lomin et al., 2015) and *Zea mays* (Yonekura-Sakakibara et al., 2004; Lomin et al., 2011; Muszynski et al., 2020).

The first step of iP- and tZ-type cytokinin biosynthesis is catalyzed by adenosine phosphate-isopentenyltransferase (IPT) to produce iP-type nucleotide precursors, iP ribotides (iPRPs) (Kakimoto, 2001; Takei et al., 2001). Then, the side chain of iPRPs is hydroxylated by cytochrome P450 monooxygenase (CYP735A) to synthesize tZ ribotides (tZRPs) (Takei et al., 2004; Kiba et al., 2013; Kiba et al., 2023). Finally, the CK-activating enzyme LONELY GUY (LOG) converts the nucleotide precursors to their active forms, iP and tZ (Kurakawa et al., 2007; Kuroha et al., 2009; Tokunaga et al., 2012). The cZ-type cytokinins are biosynthesized via the prenylation of tRNAs by tRNA-isopentenyltransferase (tRNA-IPT) and their degradation (Miyawaki et al., 2006). The iP riboside (iPR), tZ riboside (tZR) and cZ ribosides (cZR) are types of cytokinin precursors formed by the dephosphorylation of the corresponding ribotides (Sakakibara, 2006).

Active and precursor forms of cytokinins are translocated throughout plant tissues and serve as cell-to-cell and organ-to-organ signaling molecules. For instance, in the development of root vascular tissue, cytokinins produced by LOG, including LOG3 and LOG4 in xylem precursor cells, are transferred to adjacent procambial cells, thereby orchestrating cell division (Ohashi-Ito et al., 2014; De Rybel et al., 2014). In shoot apical meristems, root-borne cytokinin precursors, such as tZR, that are transported to the most apical cell layer are activated by LOG4 and LOG7 (Yadav et al., 2009; Chickarmane et al., 2012; Osugi et al., 2017). Cytokinins are then perceived by receptors expressed in inner tissues, including the organizing center (Chickarmane et al., 2012; Gruel et al., 2016; Sakakibara, 2021). On the other hand, root-borne tZ translocated via the xylem is mainly involved in leaf-size maintenance (Osugi et al., 2017). Whereas cytokinins and related molecules exhibit some degree of cell permeability, the dynamics of cytokinin flow in plants cannot be explained solely by simple diffusion. Although the basic framework of genes responsible for cytokinin biosynthesis and metabolism has mainly been elucidated (Osugi and Sakakibara, 2015; Kojima et al., 2023), genes governing cytokinin flow remain relatively unexplored.

Genes involved in several aspects of cytokinin translocation have been characterized (Zhang et al., 2023). Four types of cytokinin transporters have been reported, including PURINE PERMEASE (PUP), EQUILIBRATIVE NUCLEOSIDE TRANSPORTER (ENT), AZA-GUANINE RESISTANT (AZG), and ATP-BINDING CASSETTE (ABC) TRANSPORTER. In the PUP family, PUP1, PUP2, PUP8 and PUP14 in *Arabidopsis thaliana* and OsPUP4 in *Oryza sativa* function in the plasma membrane and play a role in transporting cytokinins (Gillissen et al., 2000; Bürkle et al., 2003; Zürcher et al., 2016; Xiao et al., 2019; Hu and Shani, 2023). Tonoplast-localized PUP7 and PUP21 can act as vacuolar cytokinin importers (Hu et al., 2023). Suppression of *PUP14* expression expanded the cytokinin signaling domain in shoot apical meristems. In contrast, the simultaneous knockdown of *PUP7*, *PUP8*, and *PUP21* narrowed the signaling domain, suggesting the involvement of *PUP*s in the regulation of apoplastic cytokinin pools to modulate perception at the plasma membrane (Zürcher et al., 2016; Hu and Shani, 2023). In the ENT family, ENT3, ENT6 and ENT8 in *A. thaliana*, and OsENT2 in *O. sativa* are thought to be localized in the plasma membrane and involved in the transport of riboside precursors (Hirose et al., 2005; Sun et al., 2005; Hirose et al., 2008). AZG1 and AZG2, members of the AZG family in *A. thaliana*, have distinct cellular localizations. AZG1 is localized solely to the plasma membrane, whereas AZG2 is found in the plasma membrane and the endoplasmic reticulum (ER) (Tessi et al., 2021; Tessi et al., 2023; Xu et al., 2024). Both proteins have been implicated in the transport of cytokinins and the regulation of root growth. In the ABC transporter family, ABCG14 in *A. thaliana* and OsABCG18 in *O. sativa* are localized to the plasma membrane and involved in long-distance transport from roots to shoots and the distribution of cytokinins and their precursors in shoots (Ko et al., 2014; Zhang et al., 2014; Zhao et al., 2019; Zhao et al., 2021a; Zhao et al., 2023). ABCG11 in *A. thaliana* has also been characterized for its involvement in modulating cytokinin responses, potentially directly or indirectly contributing to cytokinin transport (Yang et al., 2022). Furthermore, the SUGARS WILL EVENTUALLY BE EXPORTED TRANSPORTER (SWEET) HvSWEET11b in developing grains of *Hordeum vulgare* has recently been shown to transport tZ, tZR and sugars (Radchuk et al., 2023). Although these studies are informative in understanding certain aspects of cytokinin flow within plants, it is apparent that additional transporters are essential for governing cytokinin distribution.

In this study, we searched for cytokinin transport genes using a heterologous expression system and isolated a C-type ABC transporter gene, *ABCC4,* as a candidate cytokinin efflux transporter. A loss-of-function mutation and overexpression altered the root growth profile, suggesting that ABCC4 plays a role in regulating root growth and development. Additionally, larger stomatal apertures were found in loss-of-function mutants. Our findings contribute valuable insight toward understanding the intricate flow of cytokinin in plants.

## Results

### Screening of Arabidopsis genes possibly involved in cytokinin transport

To pursue genes potentially involved in cytokinin transport in Arabidopsis, we conducted a screening using the tobacco <u>s</u>yringe <u>a</u>groinfiltration and liquid chromatography-mass spectrometry (TSAL) method (Zhao et al., 2021b). In this approach, candidate genes were transiently expressed in tobacco leaf cells under the control of the cauliflower mosaic virus (CaMV) 35S promoter, followed by quantifying the concentration of cytokinins in the cellular incubation buffer. Given the abundance of transporter genes in the Arabidopsis genome, we selected genes for screening using transcriptome data reported by Yadav et al. (2009). Specifically, we chose 61 genes that were differentially expressed in domains of the shoot apical meristem and were annotated as putative plasma membrane-localized proteins and transporters with gene ontology terms (Supplementary Table S1). This screening revealed that the expression of *ABCC4* (At2g47800), a member of the C-type ABC transporter family (Kang et al., 2011), significantly enhanced the accumulation of cytokinins, namely iP, tZ and cZ, compared to the vector control (Fig. 1A). Additionally, we observed increased accumulation of the riboside and ribotide precursors, albeit to a lower extent than for the corresponding cytokinins (Supplementary Fig. S1).

**Figure 1.**
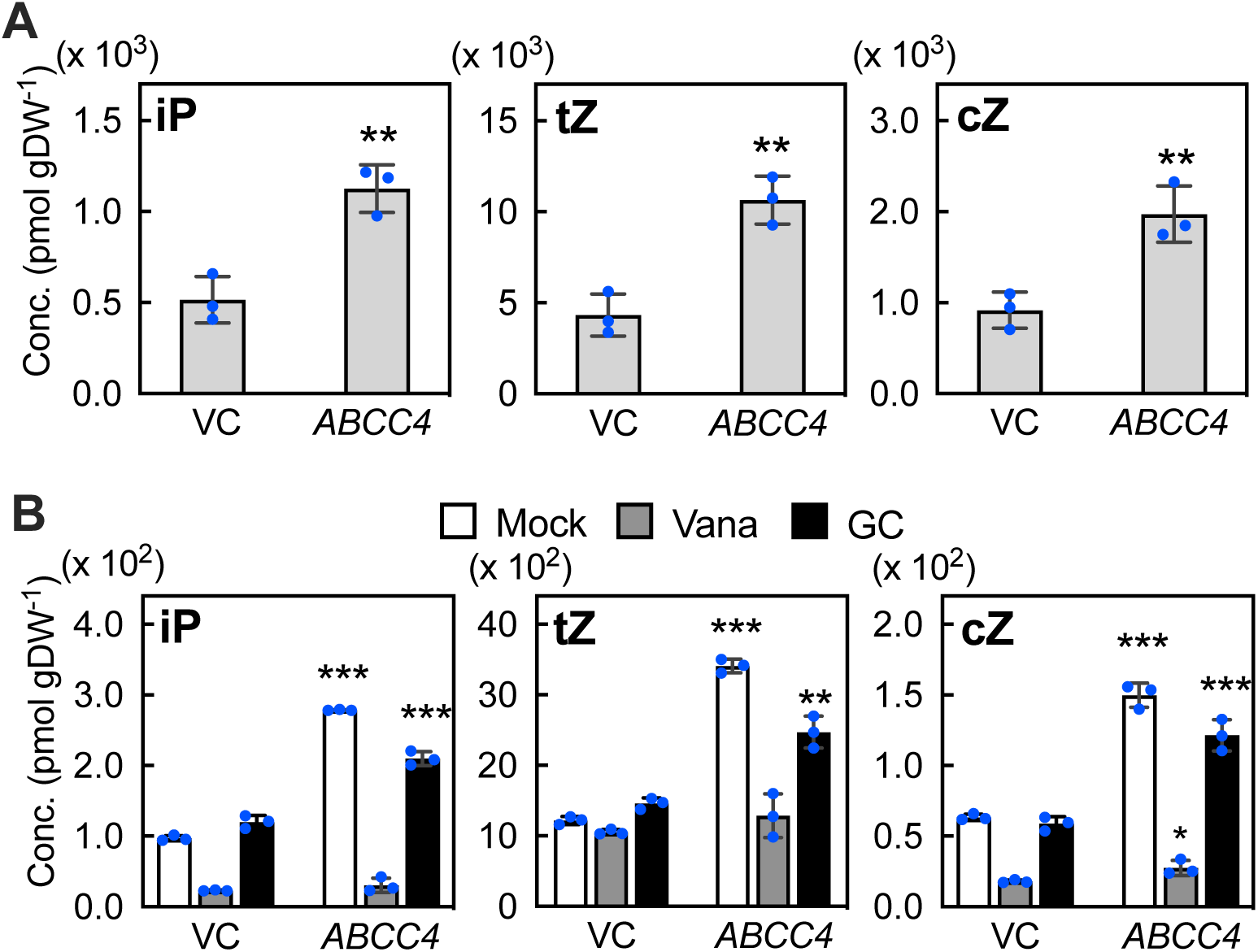
Quantification of exported cytokinins from *ABCC4*-overexpressing tobacco leaf cells. **(A)** Tobacco leaf disks expressing *ABCC4* were incubated in an incubation buffer for 12 h, followed by measurements of cytokinin levels in the buffer. Data are means ± SD (*n* = 3). Asterisks represent the Student’s *t*-test significance compared with VC (\*\**P* < 0.01). **(B)** The effect of ABC transporter inhibitors on the levels of exported cytokinins from ABCC4-overexpressing tobacco leaf cells. Tobacco leaf disks expressing *ABCC4* were incubated in incubation buffer without any inhibitors (Mock), with 1 mM orthovanadate (Vana), or with 0.1 mM glibenclamide (GC) for 12 h, after which cytokinins in the buffer were quantified. Data are means ± SD (*n* = 3). Asterisks represent the Student’s *t*-test significance compared with VC (\**P* < 0.05, \*\**P* < 0.01, \*\*\**P* < 0.001). VC, empty vector control; Conc., concentration; gDW^-1^, grams per dry weight.

To validate the involvement of ABC transporter activity in the observed effect, we employed orthovanadate, a widely used phosphate analog for inhibiting phosphatases and ABC transporters. Although the presence of orthovanadate affected the level of all cytokinins and precursors in the incubation buffer of both the vector control and *ABCC4*-expressing tobacco leaves, the inhibitor clearly diminished the enhancement accumulation of cytokinins in the *ABCC4*-expressing tobacco leaves (Fig. 1B and Supplementary Fig. S2A). In treatments with glibenclamide, an inhibitor of ABC transporters (Payen et al., 2001), the accumulation of all cytokinins was increased in the incubation buffer of *ABCC4*-expressing tobacco leaves; however, the extent of increase was distinctly reduced by the inhibitor treatment (Fig. 1B and Supplementary Fig. S2B). Collectively, these results suggest that introduced ABCC4 plays a role in the transport of cytokinins and/or their precursors in the tobacco leaf transient expression system.

### Characterization of ABCC4 as an efflux transporter of cytokinins

To further verify the role of ABCC4 in the transport of cytokinins and/or their precursors, we generated transgenic Arabidopsis lines expressing *ABCC4* under the control of a β-estradiol (βE)-inducible promoter (Zuo et al., 2000). Seedlings were incubated in MS medium in the presence and absence of βE, and the levels of cytokinins and their precursors in the medium were subsequently analyzed. In comparison to the transgenic lines without βE treatment, those with βE treatment exhibited a significant increase of iP and cZ concentrations in the medium (Fig. 2). In contrast, the levels of tZ showed an upward trend with βE treatment, the difference did not reach statistical significance. On the other hand, levels of the corresponding ribosides and ribotides either remained unchanged or decreased with βE treatment (Fig. 2).

**Figure 2.**
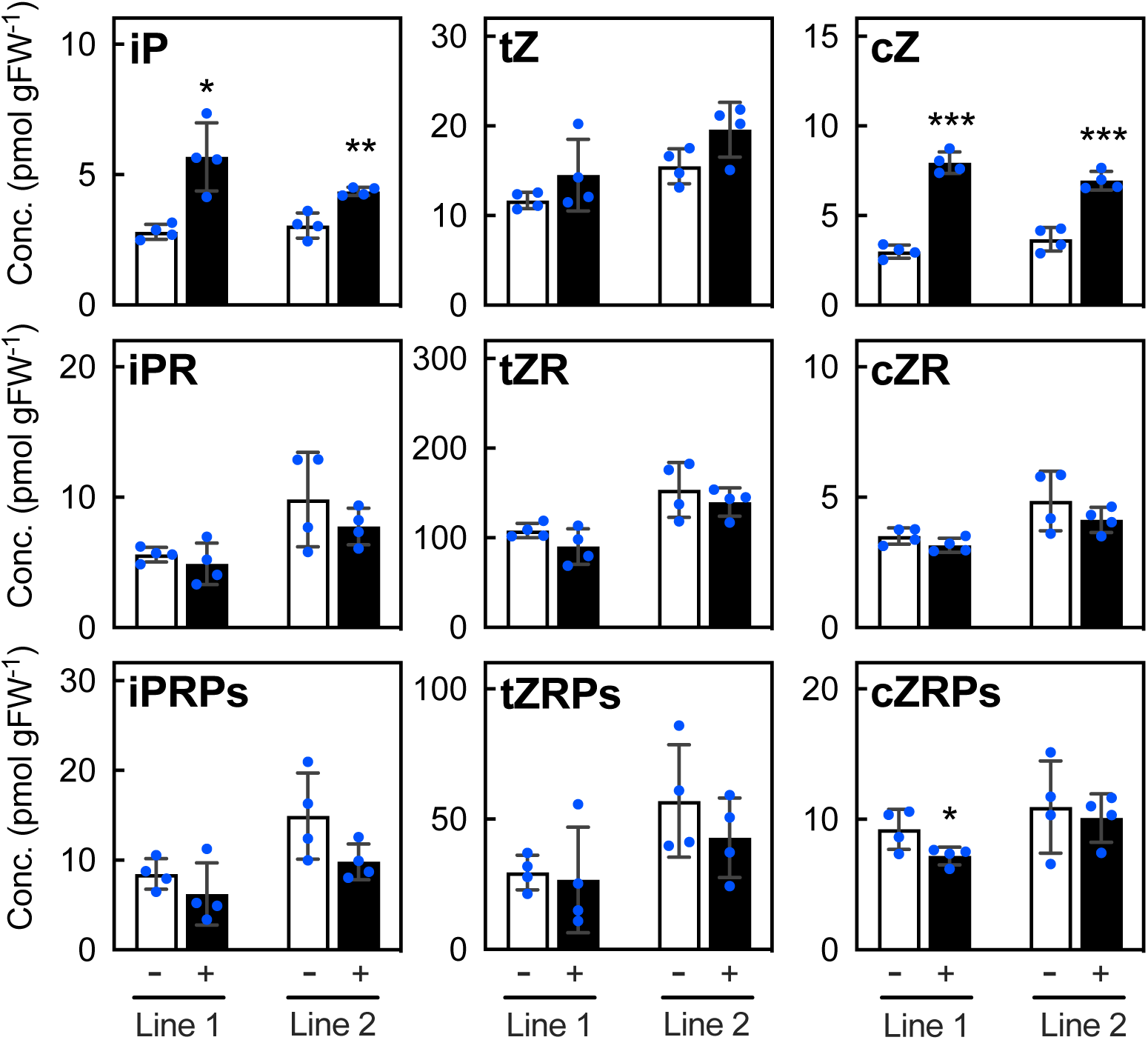
Quantification of exported cytokinins from *ABCC4*-overexpressing Arabidopsis seedlings. Seedlings of two independent transgenic Arabidopsis lines (Line 1 and Line 2) expressing *ABCC4* under control of the β-estradiol-inducible promoter were treated with (+) or without (-) 10 μM of β-estradiol for 24 h, followed by measurements of cytokinins in the culture medium. Data are means ± SD (*n* = 4). Asterisks represent the Student’s *t*-test significance compared with the mock treatment (\**P* < 0.05, \*\**P* < 0.01, \*\*\**P* < 0.001). gFW^-1^, grams per fresh weight.

In experiments using whole plants, it is difficult to determine genuine transport substrates because cytokinins can be derivatized by their metabolic enzymes. Therefore, we conducted a heterologous transport assay employing a budding yeast, *Saccharomyces cerevisiae*. In a yeast strain expressing *ABCC4* under the control of a galactose-inducible promoter, a protein band corresponding to the estimated size of ABCC4 (169 kDa) was detected (Fig. 3A). We then quantified the intracellular cytokinin levels in the yeast strains that had been fed previously with stable isotope-labeled tZ (+10) or tZR (+15), whose molecular masses exceed those of authentic tZ and tZR by 10 Da and 15 Da, respectively. As a result, the levels of tZ (+10) in yeast cells expressing ABCC4 were significantly lower in comparison to those observed in the empty vector control (Fig. 3B). In contrast, no significant difference was found in the tZR (+15) levels. These results strongly support the hypothesis that ABCC4 is involved in the efflux transport of cytokinins.

**Figure 3.**
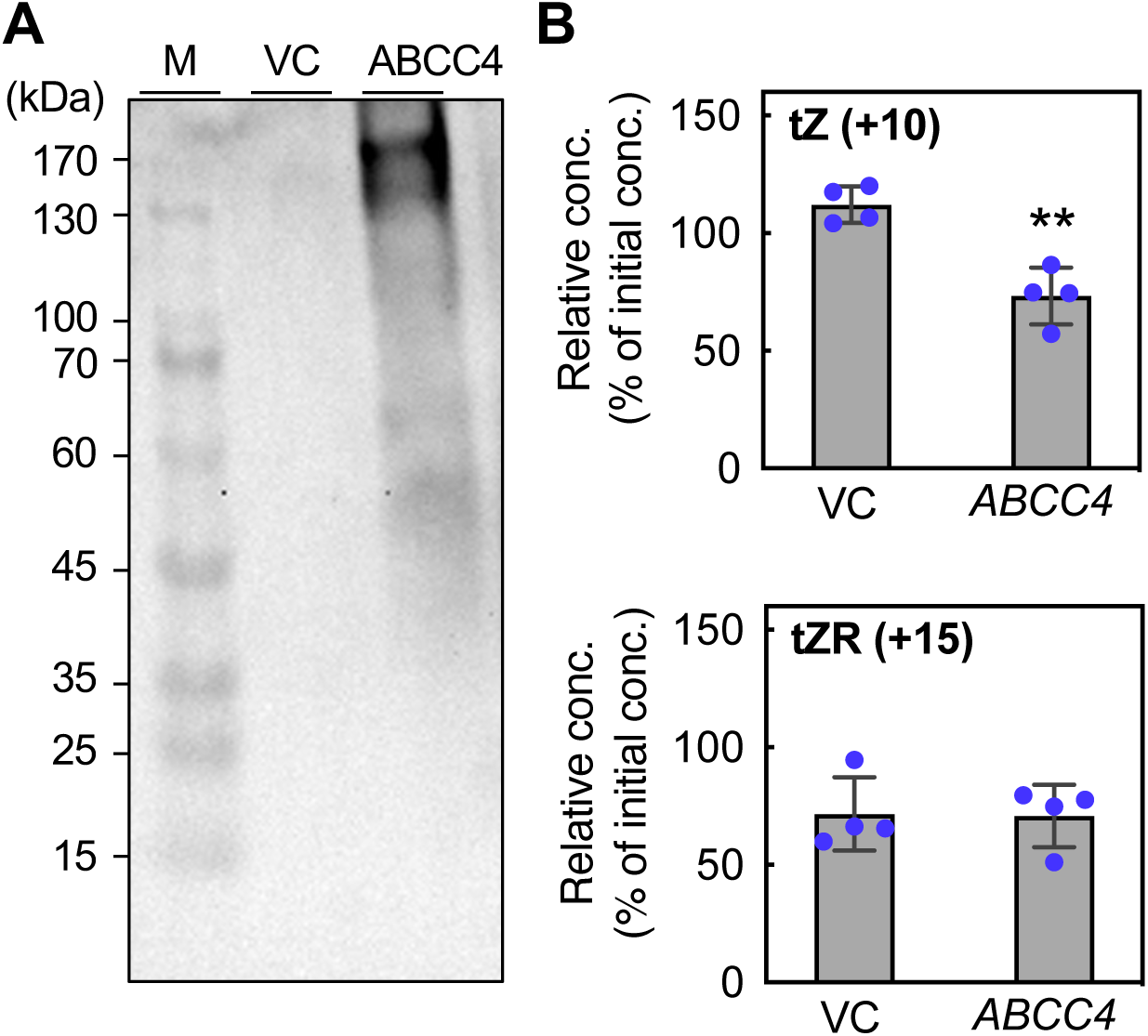
Heterologous expression of ABCC4 in yeast cells. **(A)** Immunoblot detection of ABCC4 in yeast cells. Yeast (strain YPH499) harboring the pYES-empty vector (VC) or pYES-ABCC4 (ABCC4) were cultured, and total protein was extracted. Total proteins (60 μg) were subjected to immunoblot analysis using an anti-ABCC4 antibody. Sizes of the molecular mass markers (M) are indicated on the left. **(B)** Cytokinin transport assay in yeast. VC and ABCC4 yeast strains treated with stable isotope-labeled 50 nM tZ (+10) or tZR (+15) were incubated in isotope free-buffer for 0 and 10 min, followed by quantification of the labeled compounds in the cells. The relative concentration was calculated by defining the concentration at 0 min as 100%. Data are means ± SD (*n* = 4). Asterisks represent Student’s *t*-test significance compared with VC (\*\**P* < 0.01).

### Expression analysis of *ABCC4*

ABCC4 was initially isolated as MULTIDRUG RESISTANCE-ASSOCIATED PROTEIN 4 (AtMRP4) and characterized as a transporter involved in the regulation of stomatal aperture (Klein et al., 2004). In that study, AtMPR4 was localized to the plasma membrane, and the expression was found by RT-PCR and GUS staining to be widespread throughout the plant, including the basal region of hypocotyls, primary roots, guard cells, and sepals (Klein et al., 2004). To gain more insight into the expression patterns of *ABCC4*, we conducted reverse transcription (RT)-quantitative PCR (RT-qPCR) analyses.

*ABCC4* exhibited higher expression in the roots than the shoots and shoot apices during the seedling phase and showed broad expression across all above-ground organs during the adult and reproductive phases (Fig. 4A). Since the expression of transporter genes is often regulated by its transport substrates (Jasiński et al., 2001; Swarup et al., 2008; Ko et al., 2014; Pierman et al., 2017; Zhao et al., 2019), we examined the expression of *ABCC4* in response to cytokinin treatment. Following exposure of Arabidopsis seedlings to tZ, no remarkable changes in the *ABCC4* expression were found in shoots at either 30 min or 2 h post-treatment (Fig. 4B). On the other hand, *ABCC4* expression in roots was significantly up-regulated at 30 min after cytokinin treatment, with expression levels reaching approximately three times higher than that of the mock treatment at 2 h post-treatment (Fig. 4C). These results align with the hypothesis that *ABCC4* is involved in transporting cytokinins. Indole-3-acetic acid (IAA) treatment resulted in the transient suppression of *ABCC4* expression within 30 min (Fig. 4B, C).

**Figure 4.**
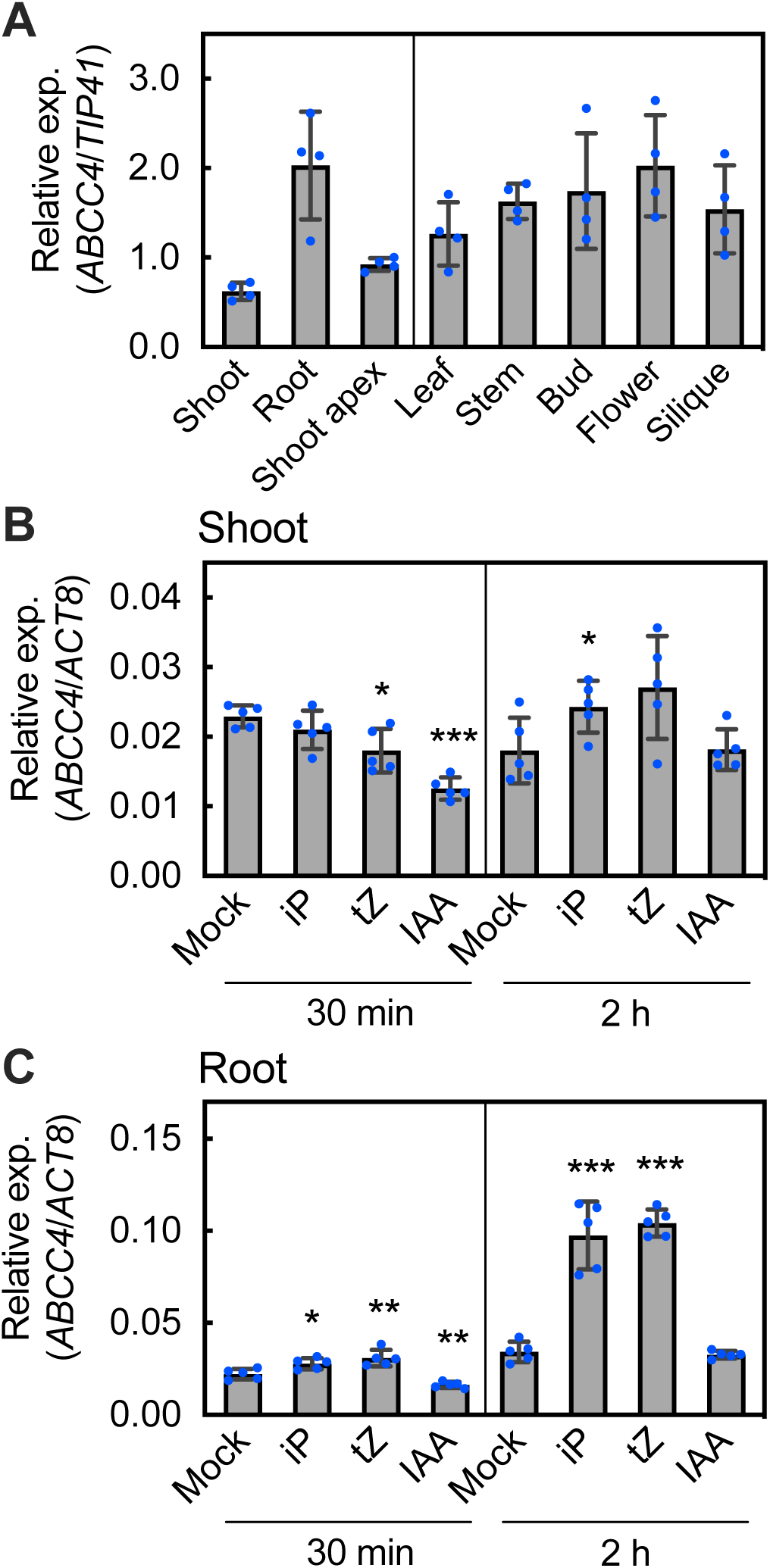
Expression patterns of *ABCC4* in Arabidopsis. **(A)** Expression patterns of *ABCC4* in plant organs. Total RNAs were extracted from shoots and roots of 10-day-old Arabidopsis seedlings (left panel) and from the indicated organs of 45-day-old plants (right panel) and subjected to RT-qPCR analysis. Expression levels of *ABCC4* were normalized to that of *TIP41,* a housekeeping gene. Data are means ± SD (*n* = 4). **(B, C)** Effect of cytokinin and auxin treatments on the expression of *ABCC4* in shoots (B) and roots (C). Arabidopsis seedlings grown for 10 d on 1/2 agar plates were sprayed with 0.01% dimethyl sulfoxide (Mock),1 μM iP (iP), 1 μM tZ (tZ), or 0.4 μM IAA (IAA). The shoots and roots were separately harvested after 30 min and 2 h. Total RNAs were extracted and subjected to RT-qPCR analysis. Expression levels of *ABCC4* were normalized to that of *ACT8.* Data are means ± SD (*n* = 5). Asterisks represent Student’s *t*-test significance compared with Mock (\**P* < 0.05, \*\**P* < 0.01, \*\*\**P* < 0.001).

### Characterization of an *abcc4* loss-of-function mutant

To explore the physiological role of *ABCC4*, we obtained a T-DNA insertional loss-of-function mutant line (*abcc4-1*) (Supplementary Fig. S3A) and generated a genome-edited frame-shift mutant line (*abcc4-2*) by the CRISPR/Cas9 mediated genome editing technique (Supplementary Fig. S4). RT-PCR analysis confirmed the absence of full-length *ABCC4* transcripts in the *abcc4-1* mutant (Supplementary Fig. S3B).

The primary roots of the loss-of-function mutants grown on 1/2MS agar plates were significantly more elongated compared to the wild type (WT) (Fig. 5A, B). Since cytokinins are known as crucial signaling molecules controlling root growth rate by determining root meristem cell number (Beemster and Baskin, 2000; Werner et al., 2003; Dello Ioio et al., 2007; Moubayidin et al., 2010), we measured the number of meristematic cells in primary roots and found that their number had increased in *abcc4* (Fig. 5C, D). On the other hand, no discernible differences were observed in lateral root growth or shoot growth when compared to the WT (Fig. 5A and Supplementary Fig. S5A, B, 6). To assess the impact of the mutation on cytokinin levels, we analyzed cytokinin concentrations in whole roots of 10-day-old *abcc4* seedlings; however, no consistent changes in the mutants compared with WT were observed (Supplementary Fig. S7A). Next, we investigated the effects of cytokinin treatment on primary root growth in the mutants.

**Figure 5.**
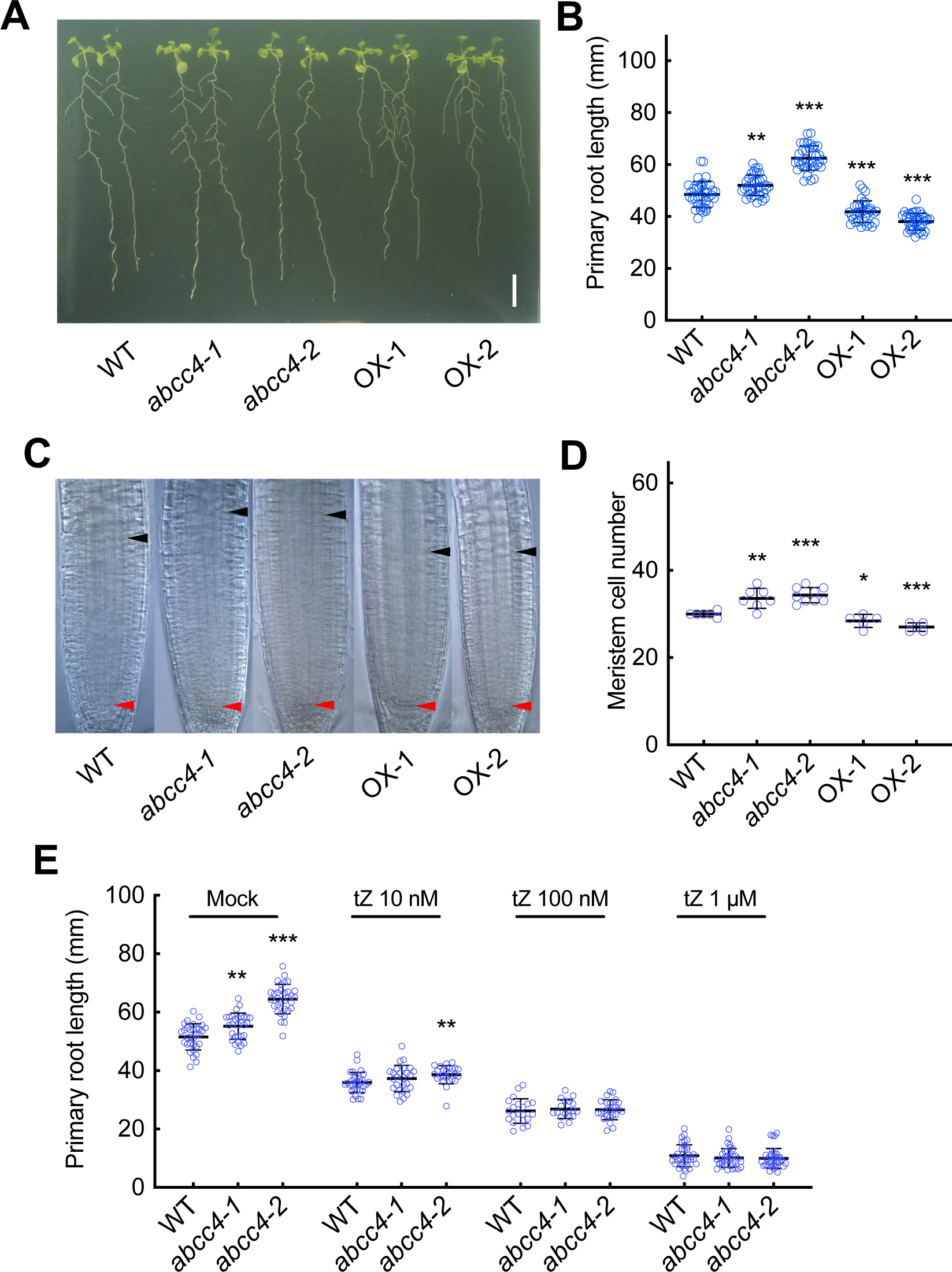
Root growth phenotypes of *abcc4* mutants and *ABCC4* overexpression lines. **(A)** A representative image of WT (Col-0), a T-DNA insertion mutant (*abcc4-1*), a genome-edited mutant (*abcc4-2*), and two *ABCC4*-overexpression lines (OX-1 and OX-2). The plants were grown on 1/2MS agar plates for 12 d. Scale bar, 1 cm. **(B)** Primary root length of WT, *abcc4-1*, *abcc4-2*, OX-1, and OX-2 seedlings grown on 1/2MS agar plates for 14 d. Data are means ± SD (*n_WT_*= 37, *n_abcc4-1_*= 34, *n_abcc4-2_*= 36, *n_OX-1_*=33, *n_OX-2_*=36). **(C)** Representative images of the root meristem of WT, *abcc4-1*, *abcc4-2*, OX-1 and OX-2 seedlings grown for 6 d. Red and black arrowheads indicate the quiescent center and the cortex transition zone, respectively. **(D)** Root meristem cell number of WT, *abcc4-1*, *abcc4-2*, OX-1 and OX-2 seedlings grown for 6 d. Data are means ± SD (*n_WT_*= 6, *n_abcc4-1_*= 7, *n_abcc4-2_*= 9, *n_OX-1_*=5, *n_OX-2_*=5). **(E)** The effect of cytokinin treatment on primary root length of WT, *abcc4-1*, and *abcc4-2.* Seedlings were grown on 1/2MS agar medium with 0.01% DMSO (Mock) or the indicated concentration of tZ for 11 days. Data are means ± SD (n = 18∼36). Asterisks represent Student’s t-test significance compared with WT (**P < 0.01, ***P < 0.001).

The length of WT primary roots declined with increasing concentrations of tZ. Primary root length of the mutants also decreased with tZ treatment, but the elongated root phenotype of the mutants was nearly abolished at 10 nM and was eliminated at higher concentrations of 100 nM and 1 µM (Fig. 5E). These results suggest that cytokinin is relevant to the primary root phenotype of the mutant.

### Analysis of phylogenetically related homologs of *ABCC4*

To investigate genes potentially sharing functions in cytokinin transport with ABCC4, we selected *ABCC14*, the phylogenetically closest homolog (Kang et al., 2011), and evaluated its cytokinin transport activity. However, no cytokinin transport capacity was detected by the TSAL method or the βE induced expression system in Arabidopsis (data not shown). We generated an *abcc4 abcc14* double mutant by the CRISPR/Cas9 system (Supplementary Fig. S4), but root and shoot growth were not different in the double mutant compared with the *abcc4* single mutant (Supplementary Fig. S8).

### Characterization of *ABCC4* overexpression lines

To gain further insights into the physiological role of *ABCC4*, we generated Arabidopsis plants overexpressing *ABCC4* driven by the CaMV 35S promoter (Supplementary Fig. S9A). No discernible morphological changes were found in the shoots of overexpressors grown in soil (Supplementary Fig. S5A, B). However, the root phenotype had fewer meristem cells, and the primary root growth of two independent *ABCC4* overexpressing lines was reduced (Fig. 5A-D). Additionally, these overexpressors exhibited higher lateral root density than the WT without altering the total density of lateral root primordia and lateral roots (Supplementary Fig. S6A, B). Furthermore, the elongation rate of lateral roots was increased (Supplementary Fig. S6C), suggesting that the emergence of lateral roots was promoted in the overexpressors.

Primary root elongation and lateral root development are reportedly regulated by auxin and cytokinin (Blilou et al., 2005; Dello Ioio et al., 2007; Laplaze et al., 2007; Marhavý et al., 2014; Du and Scheres, 2018). Therefore, we hypothesized that the levels and/or actions of auxin and/or cytokinin might be altered in the overexpressors. Consequently, we quantified IAA, cytokinins, and their precursor levels and examined the expression levels of auxin- and cytokinin-response marker genes by RT-qPCR in whole roots. Although the level of iPRPs was slightly lower, the level of IAA and all other cytokinins and their precursors in roots of ABCC4 overexpressors was not altered (Supplementary Fig. S7). Furthermore, no significant differences were detected in the expression levels of *INDOLE-3-ACETIC ACID INDUCIBLE 5* and *19* (*IAA5* and *IAA19*, respectively) and type-A *ARABIDOPSIS RESPONSE REGULATOR 5, 6,* and *15* (*ARR5, ARR6,* and *ARR15*, respectively) in overexpressors compared to the WT (Supplementary Fig. S9B and C).

### Examination of *ABCC4* involvement in root-to-shoot cytokinin transport

Given the involvement of some cytokinin transporters in organ-to-organ cytokinin transport (Ko et al., 2014; Zhang et al., 2014; Zhao et al., 2023), we conducted experiments to compare the ability of the WT, *abcc4*, and *ABCC4* overexpressors to transfer tZ from roots to shoots. The expression levels of *ARR5* in the shoots were analyzed after exogenous tZ application to the roots. As a control, we used *abcg14,* a mutant known to be impaired in root-to-shoot cytokinin transport (Ko et al., 2014; Zhang et al., 2014). The expression level of *ARR5* remained unchanged in *abcg14* upon exposure to tZ, as previously reported (Ko et al., 2014), but increased in the WT, *abcc4-1*, and *ABCC4* overexpressor (Supplementary Fig. S10). These results suggest that *ABCC4* is not essentially involved in root-to-shoot cytokinin transport.

### Effect of the *abcc4* mutation on stomatal aperture

A previous study showed that disruption of *ABCC4* (*AtMPR4*) in the Ws-2 and L*er* backgrounds resulted in larger stomatal apertures in light and dark conditions (Klein et al., 2004). In our analysis using Columbia as the genetic background, the *abcc4* mutant also showed increased stomatal apertures in the dark compared to the WT, whereas no difference was found in lighted conditions (Supplementary Fig. S11). In the overexpressors, no distinction was noted between dark and lighted conditions. This result suggests that *ABCC4* could play a role in controlling stomatal aperture.

## Discussion

In this study, we have identified and characterized *ABCC4* as a gene encoding a cytokinin efflux transporter in Arabidopsis. The cytokinin transport function was evaluated using the TSAL system, a conditional expression system in Arabidopsis, and a yeast transport assay. Identifying a cytokinin efflux transporter is noteworthy, as efflux transporters of cytokinins have been comparatively less characterized than influx transporters (Zürcher et al., 2016; Tessi et al., 2023, 2021; Hu et al., 2023). However, we were unable to examine differences in transport properties among cytokinin species, such as iP, tZ, and cZ, since competitive experiments for *in planta* exporter analysis are challenging to execute. We have attempted in vitro translation and reconstitution using liposomes (Nozawa et al., 2020) but have not yet obtained conclusive results. Other experimental systems will be required for a more in-depth analysis, including *Xenopus laevis* oocyte assays.

We found that loss-of-function mutants of *ABCC4* had elongated primary roots, whereas overexpression of *ABCC4* resulted in shortened primary roots (Fig. 5A-D). However, we could not detect any changes in the cytokinin concentration when analyzed at the whole root level (Supplementary Fig. S7A). Given that increased cytokinin action typically leads to inhibition of primary root growth (To et al., 2004; Zhang et al., 2011), a possible interpretation of our findings is that the altered action of *ABCC4* may lead to changes in local cytokinin distribution. Specifically, loss-of-function mutations in *ABCC4* may result in reduced cytokinin concentrations in the apoplast, whereas overexpression may lead to an increase in cytokinin levels. In support of this interpretation, we found that the long primary root phenotype of the mutants was abolished by external application of tZ (Fig. 5E). In addition, although cytokinins are considered to be perceived at both the ER membrane and plasma membrane (Caesar et al., 2011; Lomin et al., 2011; Wulfetange et al., 2011; Romanov et al., 2018; Antoniadi et al., 2020; Kubiasová et al., 2020), one report has suggested that cytokinin receptors are preferentially localized to the plasma membrane to perceive apoplast cytokinins in root meristematic cells (Kubiasová et al., 2020).

In addition to the primary root phenotype, the lateral root density and the lateral root elongation rate of the overexpressors were increased (Supplementary Fig. S6). Given that no alterations in lateral root growth and development were observed in the *abcc4* mutant, we assume that the lateral root phenotype of the overexpressors is irrelevant to the inherent function of *ABCC4*. Instead, the phenotype could potentially be an indirect consequence of the reduced primary root growth since it has long been known that removal or damage to the primary root results in increased lateral root formation in many plants (Thimann, 1936; Xu et al., 2017).

A previous study provided compelling evidence for the involvement of *ABCC4* (*AtMPR4*) in regulating stomatal opening (Klein et al., 2004). The loss-of-function mutants in Wassilewskija and Landsberg ecotype backgrounds had larger stomatal apertures than the wild type (Klein et al., 2004). Our analysis of *abcc4* in the Columbia background also showed enlarged stomatal apertures compared to the wild type (Supplementary Fig. S11). This finding indicates that *ABCC4* plays a role in controlling stomatal aperture across different genetic backgrounds. Cytokinins have been implicated in promoting stomatal opening (Das et al., 1976; Tanaka et al., 2006), and ABCC4 is expressed in guard cells (Klein et al., 2004), suggesting a potential relevance of the ABCC4’s cytokinin exporter activity to the stomata phenotype. Nonetheless, our current understanding of cytokinin action on stomata remains limited, necessitating further investigation in future studies.

Overall, this study revealed the multifaceted roles of *ABCC4* in plant growth and development. Additional studies are needed to elucidate the precise mechanisms underlying the effects of *ABCC4* on cytokinin flow in roots, as well as its impact on root growth and stomatal aperture. Nevertheless, our findings provide a foundation for future research to unravel the complex interplay between cytokinin distribution, perception, and growth and development.

## Materials and methods

### Plant materials and growth conditions

*Nicotiana benthamiana* (Tobacco) plants were grown in a commercial soil mixture (Supermix A, Sakata) at 23°C under long-day conditions (16-h light/8-h dark) with a photosynthetic photon flux density of 125 µmol m^-2^ s^-1^. *Arabidopsis thaliana* ecotype Columbia (Col-0) was used as the wild type. The T-DNA insertion line SALK_090215 (*abcc4-1*) was obtained from the Arabidopsis Biological Resource Center (https://abrc.osu.edu/), and its genotype was determined by genomic PCR using primers shown in Supplementary Table S2 (Supplementary Fig. S3). *Arabidopsis thaliana* seeds were sterilized and germinated on Murashige-Skoog (MS) agar plates (0.8% agar and 1% sucrose), 1/2 MS agar plates (1.1% agar and 1% sucrose), or on soil (Supermix A) at 22°C under long-day conditions (16-h light/8-h dark) with a photosynthetic photon flux density of 45 – 80 µmol m^-2^ s^-1^.

### Plasmid Construction

The coding region of *ABCC4* and *ABCC14* (with a stop codon) was amplified by RT-PCR with specific primers (Supplementary Table S2) and cloned into the pENTR/D-TOPO vector (Invitrogen) to generate an entry vector, pENTR-ABCC4. After confirmation of the sequence, the entry vector and the Gateway LR Clonase II enzyme mix (Invitrogen) were used to integrate the ABCC4 coding region into Gateway binary vectors pER8-GW-HA (Zuo et al., 2000) or pBA002-GW, a derivative of pBA002-GFP (Kiba et al., 2018), to generate pER8-ABCC4 or pBA002-ABCC4, respectively. To generate the pYES-ABCC4 plasmid for yeast transport assays, *ABCC4* was cloned into the Gateway binary vector pYES-DEST52 using pENTR-ABCC4 and Gateway LR Clonase II enzyme mix (Invitrogen).

### Agroinfiltration-based transporter activity assay in tobacco

Transient expression of genes-of-interest in tobacco leaf cells was performed according to the method described by Zhao et al. (2021b). Leaves of 30-day-old tobacco plants were infiltrated with *Agrobacterium tumefaciens* strain C58C1 harboring pBA002-ABCC4. Four days after infiltration, the leaves were cut into 3 mm × 3 mm square samples, washed twice with efflux buffer [5 mM MES-KOH buffer (pH 5.7)], and then incubated in 6 mL incubation buffer [5 mM MES-KOH buffer (pH 5.7)] at 22°C for 12 h. Aliquots of the buffer were used for cytokinin quantification. The incubated leaves were dried and weighed. For the treatment with inhibitors, either glibenclamide (final conc. 0.1 mM) or orthovanadate (final conc. 1 mM) was added to the incubation buffer.

### Cytokinin and auxin quantification

Cytokinins and auxin were semi-purified with solid-phase extraction columns and quantified using an ultra-performance liquid chromatography (UPLC)-tandem quadrupole mass spectrometer (ACQUITY UPLC System/XEVO-TQXS; Waters Corp.) with an octadecylsilyl column (ACQUITY UPLC HSS T3, 1.8 µm, 2.1 mm × 100 mm, Waters Corp.) as described (Kojima et al., 2009).

### Generation of transgenic lines

Transgenic plants were generated by the *Agrobacterium tumefaciens*-mediated floral dip method (Clough and Bent, 1998) using EHA105 strain harboring pER8-ABCC4, pER8-ABCC14, pBA002-ABCC4 or pMg137_ABCC4ABCC14. Transformants were selected on MS agar plates containing 5 μg L^−1^ bialaphos sodium salt (for pBA002) or 25 μg L^−1^ hygromycin (for pER8).

### Transporter activity assay in an Arabidopsis conditional expression system

The transporter activity assay was performed according to a previously described method (Ohyama et al., 2008). Arabidopsis seeds of β-estradiol-inducible *ABCC4* and *ABCC14* overexpression lines (T2 generation) were sown directly in 30 mL of MS liquid medium and cultured with rotation (140 rpm) under continuous light (130 µmol m^-2^ s^-1^ ) at 22°C. After 6 days, the culture medium was exchanged with new MS. After another one-day culture, transgene expression was induced with 10 μM estradiol for 24 h. Aliquots of the culture medium were subjected to cytokinin quantification.

### Synthesis of stable isotope-labeled cytokinins

tZ(+10) was synthesized from commercially available (^13^C_10_, ^15^N_5_)-adenosine 5’-monophosphate (AMP(+15)) and hydroxymethylbutenyl pyrophosphate (HMBDP) by a series of enzymatic reactions. First, AMP(+15) and HMBDP were catalyzed by recombinant Tzs (Krall et al., 2002) to form tZRMP(+15). Calf intestine alkaline phosphatase (TaKaRa) was added to the reaction mixture to generate tZR(+15). tZR(+15) was de-ribosylated by purine-nucleoside phosphorylase DeoD from *Escherichia coli* to form tZ(+10) (Takei et al., 2003). The tZ(+10) and tZR(+15) were purified using HPLC (model Alliance 2695; Waters) linked to a photodiode array detector (2996; Waters) on a reverse-phase column (SymmetryC18, 5 µm, 4.6 × 150 mm Cartridge; Waters).

### Transporter activity assay in yeast cells

pYES and pYES-ABCC4 were introduced into the YPH499 yeast strain using a Fast Yeast Transformation Kit (Geno Technology). Transformed yeast cells were grown under selective conditions in a minimal medium (46.7 g L^-1^ Minimal SD Agar Base [TaKaRa], 0.78 g L^-1^ uracil dropout supplement [TaKaRa]). Cytokinin transport assays were performed according to the method described by Zhao et al. (2023). The yeast cells were pre-cultured in a liquid yeast medium (2% raffinose) and re-suspended in an induction medium (1% raffinose and 2% galactose). After 18 h of induction, the yeast cells were incubated with an uptake buffer (100 mM potassium phosphate buffer [pH 5.8]) supplemented with 50 nM tZ (+10) or tZR (+15) at 28°C for 20 min. After washing with uptake buffer, the yeast cells were suspended and incubated in an uptake buffer for 0 and 10 min at 28°C. Cells were collected by centrifugation and subjected to cytokinin extraction and quantification. The cytokinin concentration at 0 min (initial concentration) was defined as 100%.

### Immunoblot analysis

Yeast protein was extracted by homogenizing yeast cells with acid-washed glass beads (Sigma-Aldrich) in homogenizing medium (0.25 M sorbitol, 50 mM Tris-acetate pH7.5, 2 mM EGTA-Tris, 1% PVP-40, 2mM DTT, 0.5×protease inhibitor cocktail [Sigma-Aldrich]). The protein concentration was determined using a Bio-Rad Bradford protein assay kit (Bio-Rad), and 60 μg of total protein was separated by sodium dodecyl sulfate-polyacrylamide gel electrophoresis (SDS–PAGE). The anti-ABCC4 antibodies were obtained by immunizing a rabbit with a peptide of 19 amino acid residues (Cosmo Bio) representing positions 1246–1263 (CKQFTDIPSESEWERKETL) of ABCC4. The yeast harboring pYES was used as the empty vector control.

### RT-qPCR analysis

Total RNA was extracted from plant samples using NucleoSpin RNA (Macherey-Nagel). Total RNA was used for reverse transcription by the ReverTra Ace qPCR RT Master Mix (Toyobo). Quantitative PCR (qPCR) was performed on a Quant Studio 3 Real-Time PCR system (Thermo Fisher) using the KAPA SYBR Fast qPCR kit (KAPA Biosystems) and gene-specific primer sets (Supplementary Table S3). Expression levels were normalized using *ACT8* or *TIP41* as internal controls (Kiba et al., 2018).

### CRISPR-Cas9 mutagenesis of *ABCC4* and *ABCC14*

The frame-shift mutants of *ABCC4* and/or *ABCC14* (Supplementary Fig. S4B) were generated using the transfer RNA-based-multiplex CRISPR-Cas9 vector, pMgPec12-137-2A-GFP (Hashimoto et al., 2018). Guide sequences for *ABCC4* were designed by CHOPCHOP (Labun et al., 2021), and multiplex CRISPR-Cas9 vectors pMg137_ABCC4ABCC14 harboring guide sequences for *ABCC4* and *ABCC14* were constructed as described (Hashimoto et al., 2018). pMg137_ABCC4ABCC14 contained guide sequences g4 and g14 (Supplementary Fig. S4A). Transgenic plants were generated by the *Agrobacterium tumefaciens*-mediated method using the EHA105 strain harboring the vectors. Mutations were identified by DNA sequencing of PCR products amplified with specific primer sets (Supplementary Table S2) and genomic DNA prepared from the transformants.

### Evaluation of growth phenotypes

Rosette diameters, primary root lengths, and lateral root numbers were determined from pictures using ImageJ (https://imagej.nih.gov/ij/). The stage I to stage VII primordia were counted as lateral root primordia (LRP) according to method of Malamy and Benfey (1997). LR density was calculated as the number of LR per total root length.

### Root meristem cell number analysis

The root meristem cell number for each plant was analyzed by counting the number of cortex cells in a file extending from the quiescent center to the first elongated cortex cell. Six-day-old seedlings (longer than 1 cm) viewed with a microscope (BX51; Olympus) were used to measure meristem cell numbers.

### Stomatal aperture

Stomatal aperture measurements were started in the morning after exposure to 14-16 h of darkness using plants grown for three weeks on soil. The stomatal aperture in epidermal tissues was measured as described (Hayashi et al., 2024). Leaves collected from dark-treated plants were floated on a buffer comprising 5 mM 2-ethanesulfonic acid (MES)-BTP (pH 6.5), 20 mM KCl and 0.1 mM CaCl_2_. After a light (200 µmol m^-2^ s^-1^ ) or dark treatment for 2 h, isolated epidermal fragments were prepared by blending the leaves for 3 s twice in Milli-Q water with a blender (Waring Commercial) at high speed. The epidermal fragments were collected on pieces of 58-μm nylon mesh and were immediately microphotographed using a microscope (BX50; Olympus). Stomatal apertures on the abaxial side of fragments were measured.

### Statistical analysis

All statistical analyses were conducted using Microsoft Excel. Details of the analyses are provided in the Figure legends.

## Acknowledgments

We are grateful to Drs. Fanny Bellegarde, Mimi Hashimoto, and Ryo Tabata (Nagoya University), for their helpful support and discussions.

## Author Contributions

T.U., T.Kib. and H.S. designed the experiments and analyzed the data; T.U. performed all experiments and T.Kib., Y.To, T.Kin., and H.S. supervised T.U.; Y.H. and T.Kin. helped with stomatal aperture analysis; M.K. and Y.Ta. quantified phytohormones; T.U., T.Kib. and H.S. wrote the manuscript with contributions from all co-authors; H.S. agrees to serve as the author responsible for contact and ensures communication.

## Legends to Figures

**Supplementary Figure S1. Quantification of exported cytokinin precursors from *ABCC4*-overexpressing tobacco leaf cells** Tobacco leaf disks expressing *ABCC4* were incubated in incubation buffer for 12 h, followed by measurement of cytokinin levels in the buffer. VC, empty vector control. Data are means ± SD (*n* = 3). Asterisks represent Student’s *t*-test significance compared with VC (\**P* < 0.05, \*\**P* < 0.01). gDW^-1^, grams per dry weight.

**Supplementary Figure S2. The effect of ABC transporter inhibitors on the levels of exported cytokinin precursors from ABCC4-overexpressing tobacco leaf cells** Tobacco leaf disks expressing *ABCC4* were incubated in incubation buffer in the presence of 1 mM orthovanadate (Vana), 0.1 mM glibenclamide (GC), or absence (Mock) of for 12 h, followed by measurement of cytokinin precursors in the buffer. Data are means ± SD (*n* = 3). Asterisks represent Student’s *t*-test significance compared with Mock (\**P* < 0.05, \*\**P* < 0.01, \**\*\*P* < 0.001). Crosses indicate significant differences between VC and *ABCC4* (\**P* < 0.05, \*\**P* < 0.01, \*\*\**P* < 0.001). ns, not significant. VC, empty vector control; Conc., concentration; gDW^-1^, grams per dry weight.

**Supplementary Figure S3. T-DNA insertional *abcc4-1* mutant (A)** A diagram of the T-DNA insertional site for *abcc4-1*. Boxes represent exons, and lines represent introns. The black arrowhead shows the position of the T-DNA insertion, and the arrows denote the direction and position of the PCR primers used in (B). The scale bar indicates 200 bp. **(B)** RT-PCR detection of the *ABCC4* transcripts in WT and the *abcc4-1* mutant. *ACT8* was used as the reference.

**Supplementary Figure S4. The *abcc4-2* and *abcc4 abcc14* mutants generated by the CRISPR/Cas9 system (A)** Schematic representation of CRISPR target sites. Rectangular boxes represent exons, black bars represent introns, and blue triangles identify CRISPR target sites. The bar indicates a 200 bp scale. **(B)** Partial sequences of the wild-type *ABCC4*, the wild-type *ABCC14*, the *abcc4-2* mutant, and the *abcc4 abcc14* double mutant. The sequence corresponding to a guide RNA is highlighted in yellow. Numbers represent positions in a genome sequence when the first nucleotide of the putative start codon is counted as 1. The red letter and red dashes indicate an inserted and deleted sequences, respectively. The nucleotide deletion in *ABCC4* resulted in an in-frame stop codon at the 674th amino acid. The nucleotide insertion in *ABCC14* resulted in an in-frame stop codon at the 423th amino acid.

**Supplementary Figure S5. Shoot growth phenotypes of *abcc4* mutants and *ABCC4* overexpression lines (A)** Representative images of WT, *abcc4-1*, *abcc4-2*, and two *ABCC4* overexpression lines (OX-1 and OX-2) grown for 23 d on soil. Scale bar, 1 cm. **(B)** Rosette diameter of WT, *abcc4-1*, *abcc4-2*, OX-1, and OX-2 plants grown 23 d on soil (*n = 9*).

**Supplementary Figure S6. Lateral root phenotypes of *abcc4* mutants and *ABCC4* overexpression lines (A)** Densities of lateral roots (LR) of WT, *abcc4-1*, *abcc4-2*, OX-1, and OX-2 seedlings grown for 10 d. The density was calculated by dividing the number of LR with primary root length. Data are means ± SD (*n*=11). **(B)** Total densities of LR primordia (LRP) and LR of WT, *abcc4-1*, *abcc4-2*, OX-1, and OX-2 seedlings grown for 10 d. The sum of LRP and the LR number was divided by primary root length to calculate the total density. Data are means ± SD (*n*=11). **(C)** Lateral root elongation rate of WT, *abcc4-1*, *abcc4-2*, OX-1, and OX-2 seedlings. The elongation rate was calculated by subtracting the lateral root length of 10-d-old seedlings from that of 12-d-old seedlings and dividing it by the number of days elapsed. Data are means ± SD (*n_WT_*= 18, *n_abcc4-1_*= 14, *n_abcc4-2_*= 15, *n_OX-1_*=15, *n_OX-2_*=15). Asterisks represent Student’s *t*-test significance compared with WT (\**P* < 0.05, \*\**P* < 0.01, \*\*\**P* < 0.001)

**Supplementary Figure S7. Quantification of cytokinins, cytokinin precursors, and auxin in the whole roots of the *abcc4* mutants and *ABCC4* overexpressors (A)** Quantification of cytokinins and their precursors in the whole roots of WT, *abcc4-1*, *abcc4-2*, OX-1, and OX-2 seedlings grown for 10 d on 1/2MS agar plates. **(B)** Quantification of indole 3-acetic acid (IAA) in the whole roots of WT, OX-1, and OX-2 seedlings grown for 7 d on 1/2MS agar plates. Data are means ± SD (*n* = 4∼5). Asterisks represent Student’s *t*-test significance compared with WT (\**P* < 0.05, \*\**P* < 0.01, \**\*\*P* < 0.001). gFW^-1^, grams per fresh weight.

**Supplementary Figure S8. Growth phenotype of *abcc4 abcc14* (A)** A representative image of WT, *abcc4-2*, and *abcc4 abcc14* seedlings grown for 10 d. **(B)** Primary root length of WT and *abcc4 abcc14* seedlings grown for 10 d. Data are means ± SD (*n_WT_*= 37, *n_abcc4-2_*= 36, *n_abcc4 abcc14_*=38). Asterisks represent Student’s *t*-test significance compared with WT (\*\*\**P* < 0.001) **(C)** A representative image of WT, *abcc4-2*, and *abcc4 abcc14* grown for 23 d on soil. Scale bar, 1 cm. **(D)** Rosette diameter of WT, *abcc4-2*, and *abcc4 abcc14* grown for 23 d on soil. Data are means ± SD (*n =* 8). Asterisks represent Student’s *t*-test significance compared with WT (\*\*\**P* < 0.001).

**Supplementary Figure S9. Expression levels of cytokinin and auxin-responsive marker genes in *ABCC4* overexpressing lines (A)** Expression levels of *ABCC4* in two independent *ABCC4* overexpressing lines, OX-1 and OX-2. Whole seedlings grown for 10 d were harvested. Data are means ± SD (*n* = 3). **(B)** Expression levels of auxin-response marker genes (*IAA5* and *IAA19*) in WT, OX-1, and OX-2 seedling roots grown for 10 d. **(C)** Expression levels of cytokinin-response marker genes (*ARR5, ARR6,* and *ARR15*) in WT, OX-1, and OX-2 seedling roots grown for 10 d. Data are means ± SD (*n* = 5). Expression levels were quantified by RT-qPCR analysis and normalized to *ACT8* as the internal control. Asterisks represent Student’s *t*-test significance compared with WT (\**P* < 0.05, \*\**P* < 0.01).

**Supplementary Figure S10. *ARR5* expression levels in shoots followed by tZ application in roots in an *abcc4* mutant and an *ABCC4* overexpressing line** Expression levels of *ARR5* in shoots followed by 0.01 % DMSO (Mock) or 1 μM tZ (tZ) application to roots for 30 min were analyzed in WT, *abcc4-1*, OX-1, and *abcg14* seedlings. Expression levels were quantified by RT-qPCR analysis and normalized to *ACT8* as the internal control. Data are means ± SD (*n* = 3∼4). Asterisks represent Student’s *t*-test significance compared with control of WT (\*\**P* < 0.01, \**\*\*P* < 0.001).

**Supplementary Figure S11. Stomatal aperture in WT, *abcc4* mutant and *ABCC4* overexpression lines** Measurement of stomatal aperture in WT, *abcc4-1*, OX-1 and OX-2 in the dark or light. Detached leaves were placed in dark (Dark) or lighted (Light) conditions for 2 hours. Data are means ± SD (*n* = 30). Asterisks represent Student’s *t*-test significance compared with WT (\**\*\*P* < 0.001).

